# Charting the metabolic landscape of the facultative methylotroph *Bacillus methanolicus*

**DOI:** 10.1101/858514

**Authors:** Baudoin Delépine, Marina Gil López, Marc Carnicer, Cláudia M. Vicente, Volker F. Wendisch, Stéphanie Heux

## Abstract

*Bacillus methanolicus* MGA3 is a thermotolerant and relatively fast-growing methylotroph able to secrete large quantities of glutamate and lysine. These natural characteristics make *B. methanolicus* a good candidate to become a new industrial chassis organism, especially in a methanol-based economy. This has motivated a number of omics studies of *B. methanolicus* at the genome, transcript, protein and metabolic levels. Intriguingly, the only substrates known to support *B. methanolicus* growth as sole source of carbon and energy are methanol, mannitol, and to a lesser extent glucose and arabitol. We hypothesized that comparing methylotrophic and non-methylotrophic metabolic states at the flux level would yield new insights into MGA3 metabolism. ^13^C metabolic flux analysis (^13^C-MFA) is a powerful computational method to estimate carbon flows from substrate to biomass (i.e. the *in vivo* reaction rates of the central metabolic pathways) from experimental labeling data. In this study, we designed and performed a ^13^C-MFA of the facultative methylotroph *B. methanolicus* MGA3 growing on methanol, mannitol and arabitol to compare the associated metabolic states. The results obtained validate previous findings on the methylotrophy of *B. methanolicus*, allowed us to characterize the assimilation pathway of one of the studied carbon sources, and provide a better overall understanding of this strain.

**IMPORTANCE:** Methanol is cheap, easy to transport and can be produced both from renewable and fossil resources without mobilizing arable lands. As such, it is regarded as a potential carbon source to transition toward a greener industrial chemistry. Metabolic engineering of bacteria and yeast able to efficiently consume methanol is expected to provide cell factories that will transform methanol into higher-value chemicals in the so-called methanol economy. Toward that goal, the study of natural methylotrophs such as *B. methanolicus* is critical to understand the origin of their efficient methylotrophy. This knowledge will then be leveraged to transform such natural strains into new cell factories, or to design methylotrophic capability in other strains already used by the industry.

## 1. INTRODUCTION

^13^C metabolic flux analysis (^13^C-MFA) has emerged in the last decade as an outstanding experimental method to describe the metabolic states of microorganisms. It has successfully been used to identify new pathways (1), investigate responses to environmental changes (2), improve the titer of cell factories (3), screen strains based on their enzymatic capacity (4), and more generally to provide a better understanding of the metabolism of microorganisms (5) such as methylotrophs (6–8). Briefly, ^13^C-MFA exploits a metabolic model and ^13^C-isotope patterns measured from key metabolites to estimate reaction rates consistent with the observed labeling patterns (see (9, 10) for reviews). Specifically, cells are grown on a ^13^C labeled carbon source, metabolites of the central metabolism or constituents of the biomass such as proteinogenic amino acids are sampled and quenched, and the labeling patterns of metabolites are monitored by mass spectrometry (MS) and/or nuclear magnetic resonance (NMR). With NMR carbon isotopomers are determined and hence positional information is provided whereas with MS, isotopologues are identified. A metabolic model is then used to fit these measurements to theoretical data that are simulated by optimizing reaction flux values from the metabolic model. Assuming mass balance, if the measurements are coherent and the topology of the metabolic model is correct, the experimental and simulated data will converge, yielding the estimated reaction fluxes. Finally, ^13^C-MFA provides a flux map: a predicted snapshot of the metabolic fluxes through the organism of interest during the experiment, i.e. its metabolic state. ^13^C-MFA flux maps are conceptually similar to those from flux balance analysis (FBA), a purely *in silico* method that, from a metabolic model (typically at the genome scale), computes the optimal reaction flux distribution to maximize an objective defined in terms of metabolite production, often designed to model cell growth. Importantly, ^13^C-MFA flux maps are estimates based on experimental data, whereas FBA flux maps are purely *in silico* predictions, which can be confirmed by gene deletion analysis or ^13^C-MFA for example.

*Bacillus methanolicus* MGA3 is a gram-positive bacterium that was first isolated in the 1990s from freshwater marsh soil samples after an enrichment culture on methanol at 55 °C (11). Its ability to grow quickly and to secrete large quantities of glutamate and lysine in methanol at high temperature make it a good candidate for biotech applications. Methanol is indeed viewed as a promising renewable feedstock because of its abundance and low price (12, 13), while high temperature cultures are less prone to contamination and require less cooling when scaled up (14). Furthermore, *B. methanolicus* is able to grow in seawater, which is also cheap and abundant (15). Promising metabolic engineering studies have already established MGA3 as a cell factory for the heterologous production of cadaverine (16) and GABA (17, 18). While the lack of genetic tools must have impaired the development of new applications in the past (19), the establishment of gene expression tools based on theta- and rolling-circle replicating plasmids have made *B. methanolicus* amenable to the overproduction of amino acids and their derivatives (18), and there is hope that recent breakthroughs from CRISPR interference (CRISPRi) will stimulate new innovations (20).

As a facultative methylotroph, *B. methanolicus* MGA3 can also grow on non-methylotrophic substrates such as □-mannitol, □-glucose and □-arabitol. The metabolic pathways involved in the uptake of mannitol and glucose have been described (21), as has the organization of genes involved in mannitol utilization (18). Both substrates enter the cells via a phosphotransferase system (PTS), respectively as mannitol 1-phosphate and glucose 6-phosphate, and are further converted to fructose 6-phosphate. Arabitol has recently been characterized as a fourth source of carbon and energy for *B. methanolicus* (22). The operon responsible for arabitol assimilation was identified as harboring a PTS system (AtlABC) and two putative arabitol phosphate dehydrogenases (AtlD and AtlF), whose activities were demonstrated in crude extracts. However, as the pathway was not completely characterized, it is unclear whether arabitol is assimilated through arabitol 1-phosphate to xylulose 5-phosphate (Xyl5P), or to ribulose 5-phosphate (Ribu5P) through arabitol 5-phosphate. It has been suggested that both routes operate in parallel in *Enterococcus avium* and other gram-positive bacteria (23). A series of omics studies comparing these carbon sources with methanol have contributed to a better understanding of *B. methanolicus* metabolism at the genome (21, 24), transcriptome (21, 22, 25), proteome (26) and metabolome levels (27, 28). However, a flux level description, which could validate previous findings and provide new insights into the facultative methylotrophy of *B. methanolicus* and its associated metabolic states, is still lacking.

In this study, we designed and performed ^13^C-MFA of the facultative methylotroph *Bacillus methanolicus* MGA3 growing on methanol, mannitol and arabitol. Methanol (CH_4_O) and mannitol (C_6_H_14_O_6_) are the best known carbon sources for this strain (11, 14), with comparable growth rates; while growth on arabitol (C_5_H_12_O_5_) is significantly slower (22) and its assimilation pathway has not yet been fully described. All three carbon sources are probably present in MGA3’s natural habitats, on plant leaves (29) or as plant degradation products (30). With their wide availability and fast associated growth rate, methanol and mannitol are promising feedstocks for industrial applications, while arabitol growth allows the facultative methylotrophy of MGA3 to be studied with a less efficient C source and to finish characterizing its assimilation pathway.

## 2. MATERIALS & METHODS

### 2.1 Strain

*B. methanolicus* wild-type MGA3 (ATCC 53907) strain was used for metabolic flux analyses. Strains used for cloning and expression are described in the section 2.5.1 and listed in Table 1.

**Table 1.**
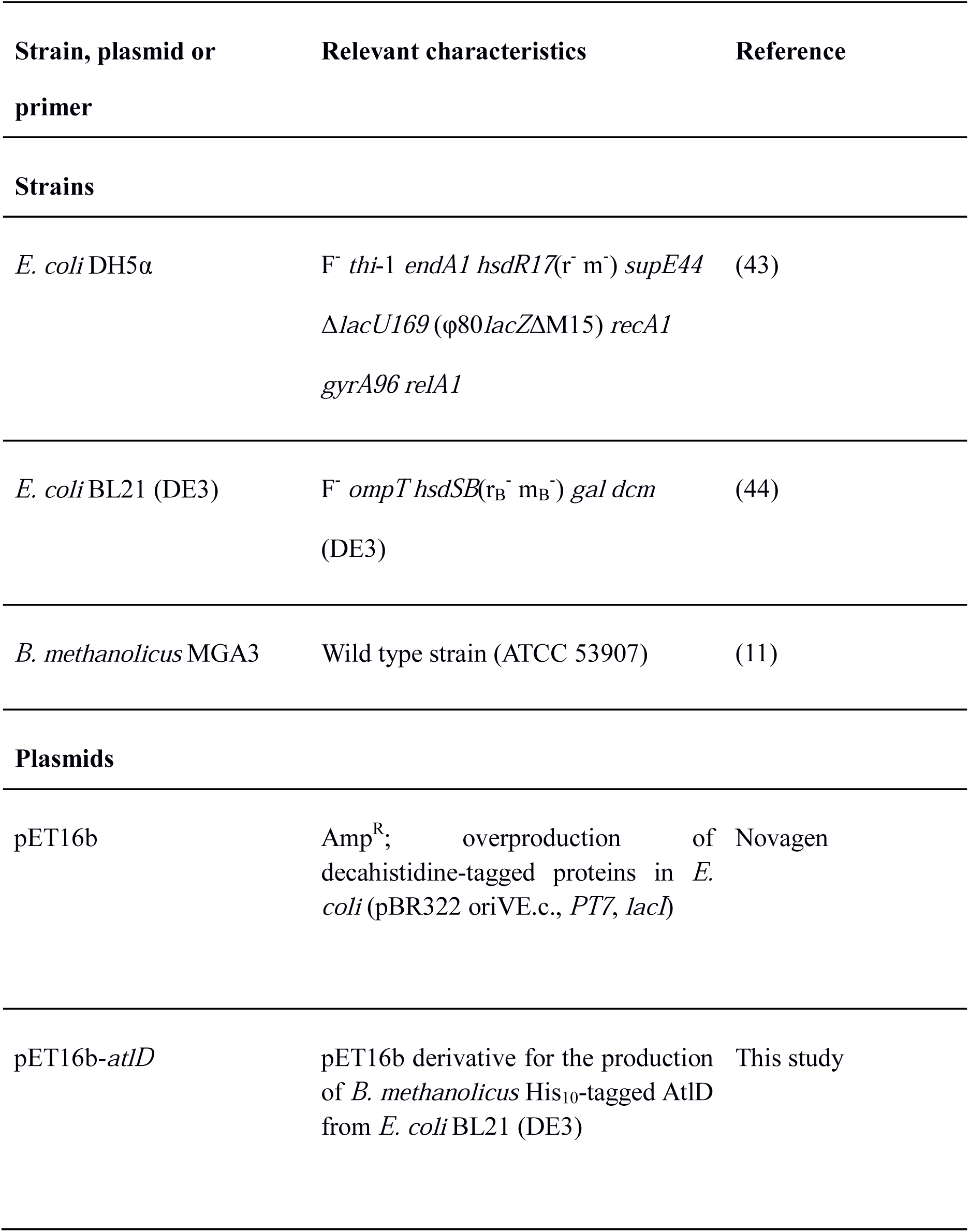
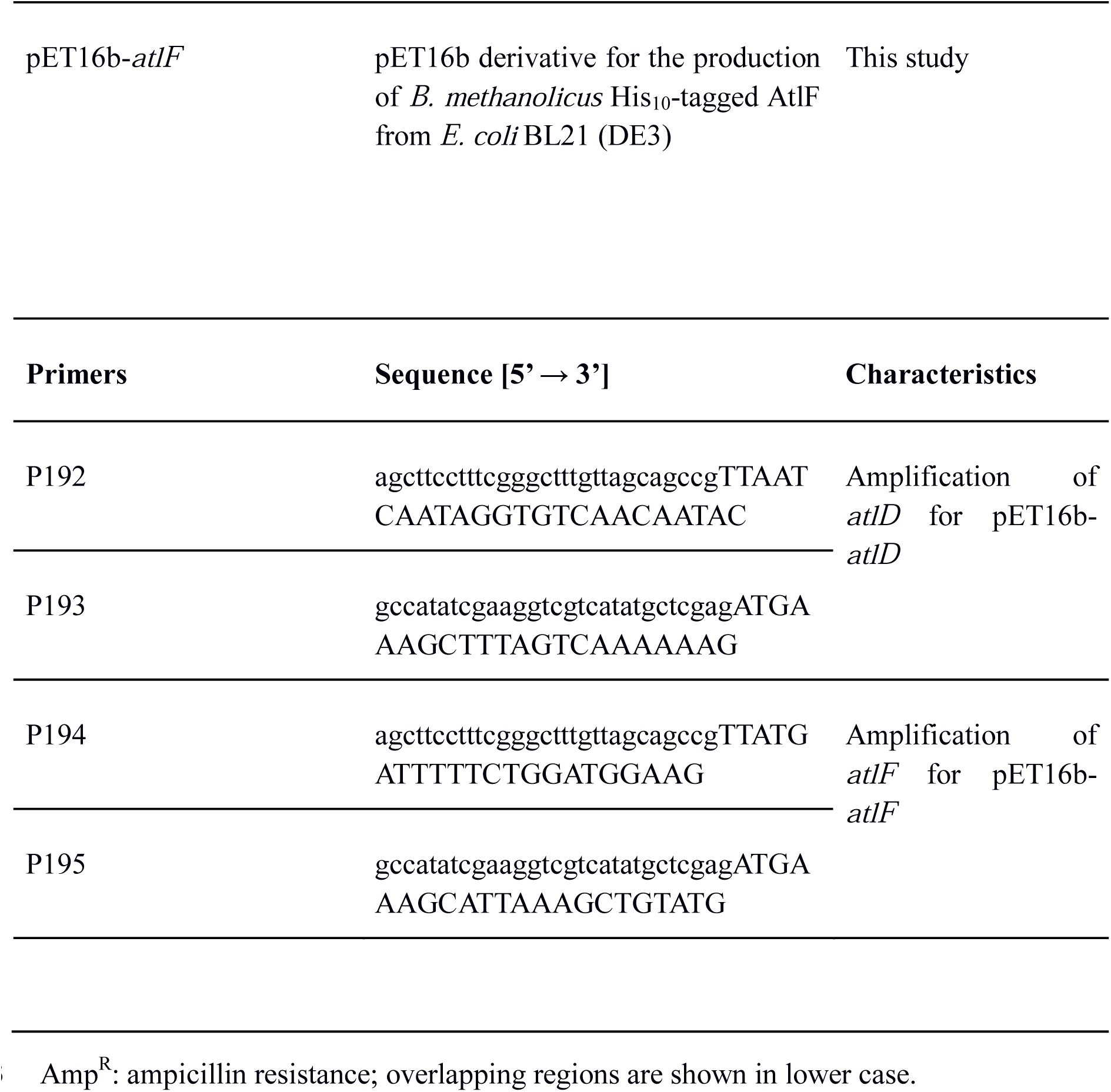
Strains, primers and plasmids used in this study.

### 2.2 Non-stationary ^13^C fluxomics experiment

#### 2.2.1 Culture conditions and parameters

For the carbon source methanol, two batch cultures were performed in 0.5 litre bioreactors (INFORS HT Multifors, The Netherlands) with a working volume of 0.40 litres coupled to a Dycor ProLine Process Mass Spectrometer (AMETEK Process Instruments, USA). The culture medium per litre was: 3.48 g Na_2_HPO_4_·12 H_2_0, 0.606 g KH_2_PO_4_, 2.5 g NH_4_Cl, 0.048 g yeast extract, 1 ml of 1 M MgSO_4_ solution, 1 ml of trace salt solution, 1 ml of vitamins solution, 0.05 ml Antifoam 204 and 150 mM of methanol. The trace salt solution per litre was: 5.56 g FeSO_4_·7 H_2_O, 0.027 g CuCl_2_·2 H_2_O, 7.35 CaCl_2_·2 H_2_O, 0.040 g CoCl_2_·6 H_2_O, 9.90 g MnCl_2_·4 H_2_O, 0.288 g ZnSO_4_·7 H_2_O and 0.031 g H_3_BO_3_. The vitamin solution per litre was: 0.10 g □-biotin, 0.10 g thiamine·HCl, 0.10 g riboflavin, 0.10 g pyridoxine·HCl, 0.10 g pantothenate, 0.10 g nicotinamide, 0.02 g p-aminobenzoic acid, 0.01 g folic acid, 0.01 g vitamin B12 and 0.01 g lipoic acid. The pre-cultures were grown in two half litre shake flasks containing 150 ml of the pre-culture medium and inoculated with cryostock of *B. methanolicus* wild-type MGA3 cells. The cultures were grown overnight at 50 °C under shaking at 200 rpm, and used to inoculate the reactors. The aeration rate of 1 vvm was controlled by a mass flow meter (INFORS HT Multifors, The Netherlands) and pO2 were maintained above 25 % throughout all cultures. Temperature, pH and stirring speed were maintained at 50 °C, pH 6.8 (with KOH 1 M) and 800 rpm, respectively. The N_2_, O_2_, Argon, CO_2_, ^13^C-CO_2_ and methanol concentrations in the bioreactors off-gas were measured on-line with the mass spectrometer.

To perform the pulse of tracer, 100 mM of ^13^C-methanol (99 % ^13^C; Euriso-Top, France) were added to the cultures at an OD_600_ of 2.5. Growth curves are available in Supplementary Data 1.

#### 2.2.2 Quantification of cells and supernatant NMR analysis

For determination of the dry weight of cells, a conversion factor of 0.389 g/l (dry weight) of cells per OD_600_ unit was used. Supernatant samples were taken to analyse substrate methanol consumption as well as by-product formation by subtracting 1 ml of culture and centrifuged it at 13000 × *g* for 60 s. Thereafter, supernatant was collected and stored until analysis at −20 °C. Supernatant analysis was performed by ^1^H 1D-NMR at 292 °K, using a 30° pulse and a relaxation delay of 20 s, with an Avance 800 MHz spectrometer (Bruker, Germany). Deuterated trimethylsilyl propionate (TSP-d4) was used as an internal standard for quantification.

#### 2.2.3 Sampling and MS analysis of intra and extracellular pool sizes

When the cultures reached an OD_600_ of 2, metabolome samples were collected using the optimized method described by (28). Briefly, total broth quenching with correction for metabolites in the extracellular medium was performed in quadruplicates to assess the metabolite pool sizes in the two cultivations performed. Metabolites pool sizes were quantified (Fig. S1) by ion chromatography tandem mass spectrometry (IC-MS/MS) using cell extract of *Escherichia coli*, cultivated on 99 % [^13^C_6_] glucose (Euroisotop, France), as internal standard (31). Liquid anion exchange chromatography was performed as described previously (32).

#### 2.2.4 Sampling and MS analysis of labeled metabolites

Label enrichments in the intracellular metabolites were followed after performing a pulse of 100 mM ^13^C-methanol at an OD_600_ 2.5. Whole broth (internal + external pools; WB) and culture filtrate (external pools; CF) were sampled to indirectly track label incorporation in the intracellular metabolites. Specifically, 13 WB and 3 CF samples were collected in 3.5 min in each bioreactor. Exact sampling times can be seen in Supplementary Data 1. IC-MS/MS quantification was used to analyse the isotopologues of each metabolite as described by (32). The metabolites analysed were PEP, Rib5P+Ribu5P+Xyl5P, Sed7P, Gnt6P, Glc6P, FruBP, Fru6P, Gly3P, 13PG, G3P+PGA, Cit, Aco, Fum, Mal and Suc. After manual peak integration, the raw peak areas were corrected for the contribution of all naturally abundant isotopes using IsoCor software (33). Some cross-contamination was found in the isotopologues M+4 of Aco and M+2 and M+3 of Gnt6P that were subsequently removed from the analysis (Supplementary Data 1).

Additionally, the exact ratio between ^13^C-methanol and ^12^C-methanol after the pulse was measured by ^1^H 1D-NMR as well as the evolution of ^12^CO_2_ and ^13^CO_2_ by the mass gas analyser.

#### 2.2.5 Mass balance

Experimental data consistency of the measured rates was verified using standard data reconciliation procedures under the elemental mass balance constraints (34, 35). The biomass elemental composition used in the reconciliation procedure was taken from the closely related non-methylotrophic bacterium *Bacillus subtilis*, CH_1.646_N_0.219_O_0.410_S_0.005_ (35, 36). The ashes content were considered to be 6 % of the dry cell weight, average value obtained from different microorganisms (i.e. *Escherichia coli, Aspergillus niger, Penicicillium chrysogenum, Klebsiella aerogenes* (37)). After no proof of mismatch in the measurements, a better estimation of the physiological parameters were obtained as described by (34).

### 2.3 Stationary ^13^C fluxomics experiments

#### 2.3.1 Culture conditions

For the carbon sources mannitol and arabitol, the culture medium composition remained unchanged. The experiments were performed in 500 ml baffled shake flasks using 40 ml of media, and cells were grown at 50 °C and 200 rpm. Pre-cultures contained yeast extract and 15 mM unlabeled mannitol or arabitol. They were inoculated as stated previously and grown overnight. The pre-cultures were used to inoculate triplicate main cultures to an OD_600_ of 0.05, after centrifugation and re-suspension in media without yeast extract. The media to perform the ^13^C MFA contained 15 mM [1-^13^C] mannitol, 15 mM [5-^13^C] arabitol or 15 mM of a mixture of 10 % [1-^13^C] arabitol and 90 % [2-^13^C] arabitol (99 % ^13^C; Omicron Biochemicals, Inc., South Bend, IN, USA). An experimentally determined conversion factor of 0.22 g/litre (dry weight) of cells per OD_600_ unit was used. Growth curves are available in Supplementary Data 1.

#### 2.3.2 Measurements of proteinogenic amino acids ^13^Cisotopologues

Mannitol (resp. arabitol) cultures were sampled around 10 h (resp. 30 h) once they reached an OD_600_ of 1.3. The pellets obtained from the cellular extract were hydrolyzed 15 h at 110 °C with 500 µl HCL 6N. Samples were evaporated and washed twice with 500 µl of ultrapure water, evaporated to dryness, resuspended (625 µl, water), and diluted (1/1000, water) for the mass spectrometry analysis.

Amino acids were separated on a PFP column (150 × 2.1 mm i.d., particle size 5 µm; Supelco Bellefonte, PEN, USA). Solvent A was 0.1 % formic acid in H_2_0 and solvent B was 0.1 % formic acid in acetonitrile at a flow rate of 250 µL/min. The gradient was adapted from the method used by (38). Solvent B was varied as follows: 0 min, 2 %; 2 min, 2 %; 10 min, 5 %; 16 min, 35 %; 20 min, 100 %; and 24 min, 100 %. The column was then equilibrated for 6 min at the initial conditions before the next sample was analyzed. The volume of injection was 20 µL.

High-resolution experiments were performed with an Ultimate 3000 HPLC system (Dionex, CA, USA) coupled to an LTQ Orbitrap Velos mass spectrometer (Thermo Fisher Scientific, Waltham, MA, USA) equipped with a heated electrospray ionization probe. MS analyses were performed in positive FTMS mode at a resolution of 60 000 (at 400 m/z) in full-scan mode, with the following source parameters: the capillary temperature was 275 °C, the source heater temperature, 250 °C, the sheath gas flow rate, 45 a.u. (arbitrary unit), the auxiliary gas flow rate, 20 a.u., the S-Lens RF level, 40 %, and the source voltage, 5 kV. Isotopic clusters were determined by extracting the exact mass of all isotopologues, with a tolerance of 5 ppm. Experimental carbon isotopologue distributions (CIDs) of alanine, glycine, valine, serine, threonine, phenylalanine, aspartate, glutamate, histidine, isoleucine, leucine, lysine, arginine, tyrosine, proline and methionine were obtained after correction of raw MS data for naturally occurring isotopes other than carbon, using IsoCor (33).

Careful inspection of the CIDs revealed an overall excellent reproducibility between both the technical and biological replicates (Supplementary Data 1). However, M+0 of valine and M+0-M+1 of glycine had a higher variability which could be due to a signal closer to the noise level.

#### 2.3.3 NMR measurements

Concentrations in supernatants were measured by ^1^H 1D-NMR at 290 °K, using a 30° angle pulse and a presaturation of water signal was applied during a relaxation delay of 8 s. TSP-d4 was used as internal standard for calibration and quantification.

The measurement of isotopomers and specific enrichments of targeted biomass components were performed using the same samples used for proteinogenic amino acids ^13^C-isotopologues MS analysis, redried and suspended in 200 μl D_2_O (0.1 % DCl). The positional isotopomer distribution of alanine C2 and C3 was extracted from the analysis of ^13^C–^13^C couplings in 2D ^1^H–^13^C HSQC experiments (39). The carbon isotopic enrichments of alanine (C2 and C3) and histidine (C2 and C5) were extracted from the analysis of ^1^H–^13^C couplings using 2D-zero quantum filtered-TOCSY (ZQF-TOCSY) (40).

NMR spectra were collected on an Avance III 800 MHz spectrometer (Bruker, Germany), equipped with a 5mm z-gradient QPCI cryoprobe. Every acquisition 1D, 2D and absolute quantification were performed on ^1^H-1D spectra using TopSpin 3.5 (Bruker, Germany).

### 2.4 Models and simulations

#### 2.4.1 Metabolic Flux Analysis software

All simulations needed for Metabolic Flux Analysis (MFA) on methanol, mannitol and arabitol were performed with *influx_si* software v4.4.4 (41), either in stationary or non-stationary mode. *influx* has the advantage to allow for the integration of labeling data coming from different experimental setups (MS, NMR) and to support several integration strategies (stationary, non-stationary, parallel labeling).

All input and output files needed for reproducibility are available in an archive in Supplementary Data 2. Importantly, this includes the models in FTBL and SBML file formats.

#### 2.4.2 Metabolic models

We designed the models to cover central carbon metabolism and biomass needs of *B. methanolicus*. *influx_si* uses a non-standard legacy file format (FTBL) to encode the metabolic network and the associated atom-atom transitions. This format centralizes the metabolic network with all biological measurements, i.e. metabolites pool sizes, fluxes, and carbon isotopologue distribution. Consequently, we developed distinct model files for each carbon source. Models share the same nomenclature and the same general topology between them, which is displayed in Fig. 1 and detailed in Supplementary Data 1.

**Fig. 1:**
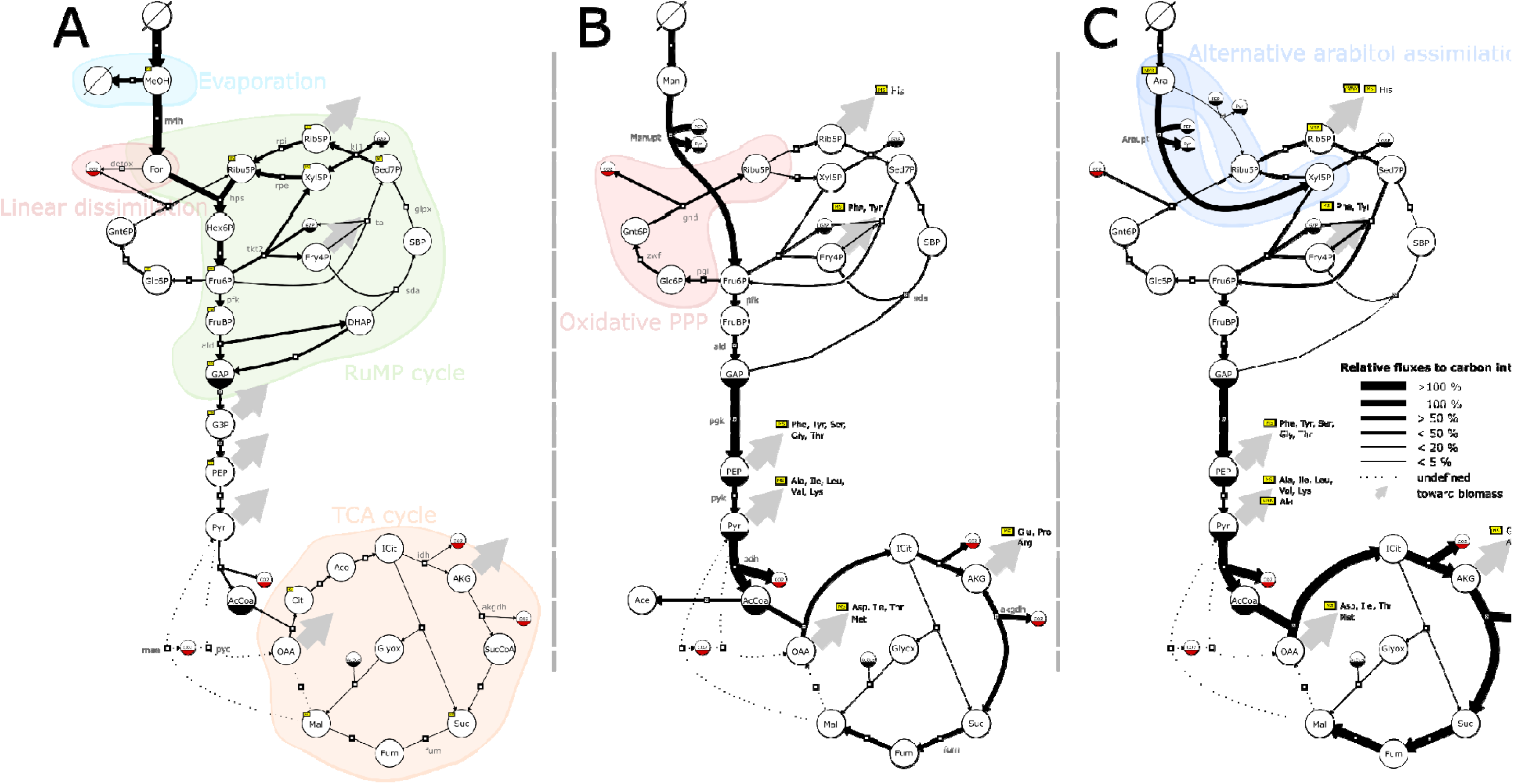
Flux map of *B. methanolicus* central metabolism relative to substrate intake for methanol (A), mannitol (B) and arabitol (C). The thickness of the reaction lines represent the flux proportions relative to substrate incorporation into the metabolism: hps (A), Manupt (B), Araupt (C). Flux proportions were averaged among the biological replicates (n_methanol_=2, n_arabitol_ = n_mannitol_ = 3). The direction of the reaction arrows represent the estimated net flux directionality. Large grey arrows represent a flux directed toward amino acid synthesis and biomass requirements for growth. Pathways discussed in the text are represented by a background patch of color. Metabolites are represented by circles and have a solid bottom half if they are duplicated on the same panel, as per recommended by the SBGN standard. Metabolites subjected to experimental measurement of ^13^C labelling are marked by a yellow box. Some intermediate metabolites are not needed for ^13^C-MFA and are consequently lumped in mannitol and arabitol models (B, C); the same metabolites can be explicitly modelled in methanol model (A) since non-stationary ^13^C-MFA requires all intermediates pools (e.g.: G3P, Cit, Aco). Reactions toward amino acid pools and biomass are not represented.

Assimilation pathways of methanol, mannitol and arabitol were included when relevant to explain the incorporation of the tracer. The CO_2_ pool was explicitly modeled within the system to allow for re-incorporation of tracer via CO_2_. Amino acids synthesis pathways were modeled as part of biomass needs along with important precursors. Biomass equation was borrowed from *Bacillus subtilis* genome-scale model (42). Reaction for phosphoenolpyruvate carboxykinase (BMMGA3_RS13120) was not included in the final model as the associated expression level was rather low in proteomics (26) and transcriptomics studies (21, 25).

Unless otherwise stated, each culture replicate was processed independently to estimate fluxes for their respective carbon source condition.

For MFA on mannitol, we exploited carbon isotopologue distribution data of proteinogenic alanine, glycine, valine, serine, threonine, phenylalanine, aspartate, glutamate, histidine, isoleucine, leucine, lysine, arginine, tyrosine, proline and methionine. Acetate production was modeled with an export flux from acetyl-CoA to which we associated the acetate flux measured from supernatant data. Acetyl-CoA was further constrained using acetate positional labeling measured by ^1^H 1D-NMR spectroscopy from the supernatant.

For MFA on arabitol, we averaged the proteinogenic labeling measurements of the [5-^13^C] arabitol experiment and exploited it as a parallel labeling dataset for each biological replicate of the [1/2-^13^C] arabitol experiment. We analysed the same proteinogenic amino acids as those mentioned above for mannitol. No acetate was observed in the supernatant and the associated export reaction was consequently excluded from this model. Additionally, we exploited specific labeling enrichment of histidine, alanine and ribulose-5-P, and positional isotopomer data of alanine from labeling samples as described in section 2.3.3.

For MFA on methanol, the non-stationary nature of the experiment and the subsequent importance of the pools on flux distributions forced us to explicit most of the reactions of the central metabolism that were lumped for the stationary models. Measurements described above were used to constrain intra and extracellular pools (section 2.2.3), and isotopologues profiles (section 2.2.4) through *influx_si* optimization process. Only the measurement of the CO2 from the exhaust gas was used but not it’s labelling dynamic. The exchange fluxes of CO_2_ and methanol (feed and evaporation) were also exploited. No acetate production was observed. Because some isotopologues of Gnt6P (M+2 & M+3) could not be quantified, these isotopic measurements could not be included in the *influx_si* model, hence the flux through the oxidative PPP could not be accurately resolved. To exploit these partial data and estimate the flux through the oxidative PPP, we applied the ScalaFlux approach (43), which allows to quantify fluxes through a given metabolic subnetwork of interest by modeling label propagation directly from the metabolic precursor of this subnetwork. The flux calculations are thus purely based on information from within the subnetwork of interest, and no additional knowledge about the surrounding network is required. Briefly, we defined a metabolic subnetwork containing two reactions: i) one reaction converting Glc6P into Gnt6P, and ii) one sink reaction operating at the same rate to balance the 6PG pool, consistently with the metabolic steady-state assumption. This metabolic model assumes direct formation of Gnt6P from Glc6P. Flux calculations were based on the time-course profile of M+0 (relative to M+0,M+1,M+4,M+5,M+6), which showed the highest range of variation during this experiment and was thus less affected by measurements noise. Fraction M+0 of Glc6P was corrected for the presence of an extracellular unlabeled pool by normalizing its profile to the isotopic steady-state of Gnt6P. Experimental labelling dynamics of the precursor of this subnetwork (Glc6P) were used as label input, and the flux through the oxidative PPP was estimated by fitting the concentration and M+0 profile of Gnt6P. The goodness of fit was evaluated using a chi-square test, and the flux precision was estimated using the Monte-Carlo approach (with 200 iterations). Using such approach we obtain a flux through the oxidative PPP (i.e. zwf) of: 0.5909 +-0.0578 and 0.6919 +-0.0959 µmol/gDCW/s for the two replicates. Next we used those data to constraint the flux through zwf (as a “measured flux”) through *influx_si* optimization process.

#### 2.4.3. Quality checks

Experimental data were fitted to our models as per described above. For each culture replicate we performed a Monte Carlo sensitivity analysis (n=100) on the fit to assess its robustness to small variations around the fitted values. We also performed a chi-squared goodness-of-fit statistical test to ensure that simulated data for each biological replicate were significantly close to experimental data. All tests were significant with a significance level (□) of 0.05 (Supplementary Data 2). For convenience, we provide figures of measured vs. simulated data points (Fig. S3).

Let us note here that we unfortunately were unable to estimate at a satisfactory precision the fluxes through malate dehydrogenase (BMMGA3_RS12590, 1.1.1.37), malic enzyme (*mae*, BMMGA3_RS12650, 1.1.1.38 or 1.1.1.40) and pyruvate carboxylase (*pyc*, BMMGA3_RS05255, 6.4.1.1). Those three reactions formed a cycle from pyruvate (Pyr) to oxaloacetate (OAA) and malate (Mal) in all our tested models. However, the Monte Carlo sensitivity analysis revealed that those fluxes were statistically undefined (not shown), meaning that virtually any value through the cycle would satisfy the constraints of the rest of the network. In the absence of biological data to motivate any new constraint on the model, we preferred to leave those reactions out of the analysis.

### 2.5 Analysis of arabitol phosphate dehydrogenases AtlD and AtlF

#### 2.5.1 Strains and culture conditions

In this study, *Escherichia coli* DH5α (44) was used as the standard cloning host and recombinant protein production was carried out with *E. coli* BL21(DE3) (45). A summary of the strains, primers and plasmids constructed and used in this study can be found in Table 1. *E. coli* strains were routinely cultivated at 37 °C and 180 rpm in Lysogeny Broth (LB) medium or on LB agar plates supplemented with 100 µg ml^−1^ ampicillin and 0.5 mM IPTG when relevant.

#### 2.5.1 Recombinant DNA work

Molecular cloning was performed as previously described (46) using primer sequences listed in Table 1. Total DNA isolation from *B. methanolicus* was performed as described in (47). Inserts were amplified by polymerase chain reactions (PCRs) with ALLin^TM^ HiFi DNA Polymerase (HighQu, Kraichtal, Germany) and purified with the NucleoSpin® Gel and PCR Clean-up kit (Macherey-Nagel, Düren, Germany). Plasmids were constructed from PCR-generated fragments and pET16b vector cut with restriction enzymes using the isothermal DNA assembly method (48). The GeneJET Plasmid Miniprep Kit (Thermo Fisher Scientific, Waltham, USA) was used for plasmid isolation. For the transformation of chemically competent *E. coli* cells, the procedure described by (49) was followed. Colony PCRs were performed using Taq polymerase (New England Biolabs, Ipswich, England) with primers P192, P193, P194 and P195 (Table 1). All cloned DNA fragments were verified by sequencing (Sequencing Core Facility, Bielefeld University).

#### 2.5.2 Overproduction and purification of AtlD and AtlF

Plasmids for protein production using *E. coli* BL21 (DE3) were constructed on the basis of pET16b (Novagen, Madison, WI, USA) and are presented in Table 1. The *atlD* and *atlF* genes were PCR-amplified from *B. methanolicus* MGA3 genomic DNA using the primers P192 and P193 or P194 and P195, respectively (Table 1). The resulting product was joined with *Bam*HI digested pET16b by applying the isothermal DNA assembly method (48), resulting in pET16b-*atlD* and pET16b-*atlF*. The pET16 vector allows for production of N-terminal His_10_-tagged proteins. Protein production and purification was performed following the indications of (50), except for cell lysis which was performed by sonication (UP 200 S, Dr. Hielscher GmbH, Teltow, Germany) on ice at an amplitude of 55 % and a duty cycle of 0.5 for 8 min with a pause in between. Supernatants were subsequently filtered using a 0.2 µm filter and purified by nickel affinity chromatography with nickel-activated nitrilotriacetic acid-agarose (Ni-NTA) (Novagen, San Diego, CA, USA). His-tagged AltD and AtlF proteins eluted with 20 mM Tris, 300 mM NaCl, 5 % (vol/vol) glycerol and 50, 100, 200, or 400 mM imidazole were analysed by 12 % SDS-PAGE (51). Fractions showing the highest protein concentrations (with 100 and 200 mM or 100, 200 and 400 mM imidazole for AtlD and AtlF, respectively) were pooled and protein concentration was measured according to the Bradford method (52) using bovine serum albumin as reference. The purified protein was subsequently applied for enzymatic assays.

#### 2.5.3 Arabitol phosphate dehydrogenase enzymatic assays

Determination of purified AtlD and AtlF activities in the reductive reaction using Xyl5P or Ribu5P as substrate were performed as previously described (23). The assay mixture contained 20 mM Tris-HCl buffer (pH 7.2), 1 mM DTT, 0.04 to 0.3 mM NADH or NADPH, 0.03 to 0.6 mM Xyl5P or 0.2 to 4 mM Ribu5P and 0.01 to 0.04 mg AtlD or 0.2 to 0.4 mg AtlF in a total volume of 1 ml. The oxidation rate of NADH or NADPH was monitored at 340 nm and 30 °C for 3 min using a Shimadzu UV-1202 spectrophotometer (Shimadzu, Duisburg, Germany). In order to confirm the presence of arabitol phosphate in the enzyme reactions after reduction of Xyl5P and Ribu5P, samples were subjected to liquid chromatography-mass spectrometry (LC-MS) analyses following the procedure described in (53).

### 2.6 Data availability

Gene locii mentioned throughout the text are from NCBI annotation of *B. methanolicus* MGA3 genome NZ_CP007739.1 (https://www.ncbi.nlm.nih.gov/nuccore/NZ_CP007739.1).

Supplementary Data 1 contains growth curves, processed MS and NMR data, a summary of the reactions modelled and the absolute fluxes in mmol/gDCW/h.

Supplementary Data 2 contains raw models, input and output files for influx software and can be downloaded from https://fairdomhub.org/data_files/3269?version=2.

## 3. RESULTS & DISCUSSION

### 3.1 *In vitro* assessment of arabitol assimilation

The operon responsible for arabitol assimilation in *B. methanolicus* consists of a PTS system (AtlABC) and two putative arabitol phosphate dehydrogenases (AtlD and AtlF) (22), which are chromosomally encoded and belong to the diverse superfamily of medium-chain dehydrogenases/reductases (MDRs). However, the physiological roles of AtlD and AtlF have not been described to date. Members of the MDR superfamily have high sequence conservation, but sequence-based prediction of their substrate scope is difficult since wide substrate specificity is common (54, 55) (see Supplementary Text for a discussion of their phylogeny). The substrate selectivity of AtlD and AtlF was therefore studied to identify whether arabitol is assimilated via arabitol 1-phosphate dehydrogenase to Xyl5P, and/or to Ribu5P via arabitol 5-phosphate dehydrogenase (23) (Fig. 2). The enzymes were purified as N-terminally His-tagged proteins from recombinant *E. coli* by nickel chelate chromatography. Arabitol phosphate oxidation could not be assayed because there are no commercial arabitol 1-phosphate or arabitol 5-phosphate standards. Therefore, the reverse reaction was tested, as described in material and methods, with either Xyl5P or Ribu5P as substrate for NAD(P)H dependent reduction. A number of potential sugars (□-ribose, □-fructose, □-xylose, □-mannose, □-arabinose, □-arabinose, □-sorbose, □-galactose, □-glucose, ribose 5-phosphate, glucose 6-phosphate, glucose 1-phosphate, fructose 6-phosphate and fructose 1-phosphate) and sugar alcohols (□-arabitol, □-mannitol, □-galactitol, □-sorbitol, □-arabitol, □-maltitol, □-xylitol, ribitol and meso-Erythritol) were also tested as substrates but no significant activity (i.e. > 0.05 U mg^−1^) was detected. The kinetic parameters measured for AtlD and AtlF reduction using either Xyl5P or Ribu5P as substrate are summarized in Table 2. The *K*_M_ of 0.07 ± 0.03 mM for AtlD with Xyl5P as substrate and NADH as cofactor was 2.5 times lower than the value obtained for AtlF (0.18 ± 0.05 mM). With Ribu5P as substrate, the *K*_M_ for AtlD was 17 times lower (1.21 ± 0.42 mM) (Table 2), and for AtlF, no activity was detected. The *V*_max_ for AtlD with Xyl5P as substrate and NADH as cofactor was 2.7 times higher than with Ribu5P as substrate (0.49 ± 0.06 U mg^−1^) and 12 times higher than for AtlF with Xyl5P (0.11 ± 0.01 U mg^−1^) (Table 2). NADH was the preferred cofactor over NADPH with a ten times lower *K*_M_ (0.01 ± 0.01 vs 0.11 ± 0.09 mM). The affinity and NADPH dependent activity (0.24 ± 0.06 U mg^−1^) could only be determined with AtlD and Xyl5P, as no significant activity was detected for any of the other reactions. LC-MS analyses were performed to confirm the formation of arabitol phosphate in the enzyme reactions catalyzed by AtlD. Although arabitol 1-phosphate and arabitol 5-phosphate could not be distinguished, arabitol phosphate was clearly produced with both Xyl5P and Ribu5P as substrates (Fig. S2).

**Fig. 2:**
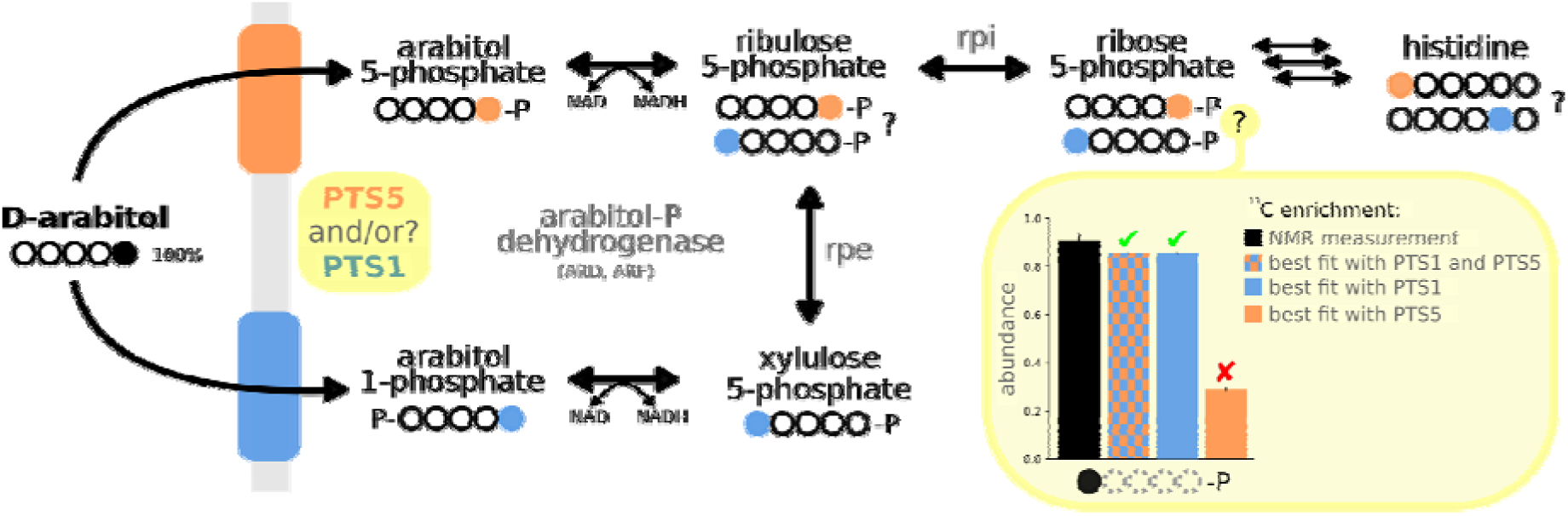
Alternative assimilation pathways of D-arabitol with an example of the measured and expected labeling of Rib5P. Arabitol entry point into the metabolism is expected to be Ribu5P or Xyl5P, depending on the substrate specificity of the PTS system and the arabitol phosphate dehydrogenase. Taking [5-^13^C]arabitol as an example, we show that the labeling of downstream metabolites can be used to identify which pathway is operating *in vivo* (Ribu5P, orange; Xyl5P, blue). We exemplify our approach with one data point: the NMR specific enrichment of Rib5P and the associated estimates from the best fit of different scenarios in which PTS1 or PTS5 are allowed to carry flux in the model (barplot). Unlike the others, the model in which only PTS5 was allowed to carry flux did not fit the data (red crossmark, □). Circles are carbon atoms: solid for ^13^C (●), empty for ^12^C (○), dotted when irrelevant.

**Table 2.** Kinetic data of purified AtlD and AtlF

| Condition<sup>a</sup>: protein,<br>substrate, cofactor<sup>b</sup> | Substrate $K_M$ (mM) | $V_{max}$ (U mg<sup>-1</sup>)<sup>c</sup> | $V_{max} / K_M$ |
| --- | --- | --- | --- |
| AtlD, Xyl5P, NADH | 0.07 ± 0.03 | 1.33 ± 0.23 | 19 |
| AtlD, Ribu5P, NADH | 1.21 ± 0.42 | 0.49 ± 0.06 | 0.4 |
| AtlF, Xyl5P, NADH | 0.18 ± 0.05 | 0.11 ± 0.01 | 0.6 |

<sup>a</sup> The following reactions were also analysed, but no significant activity (i.e. < 0.05 U mg<sup>-1</sup>) could be detected: AtlD, Ribu5P, NADPH; AtlF, Xyl5P, NADPH; AtlF, Ribu5P, NADH; AtlF, Ribu5P, NADPH.
<sup>b</sup> Cofactor $K_M$ values were analysed using 0.2 mM Xyl5P and were 0.01 ± 0.01 and 0.11 ± 0.09 for NADH and NADPH, respectively.
<sup>c</sup> The $V_{max}$ for AtlD, Xyl5P, NADPH was calculated using 0.2 mM of substrate and 0.3 mM of cofactor, and resulted in 0.24 ± 0.06 U mg<sup>-1</sup>.

Overall, these data suggest that AtlD has a major role in arabitol catabolism *in vitro* and that AtlD and AltF both prefer Xyl5P. This suggests that arabitol is mainly assimilated through the arabitol 1-phosphate pathway. However, assimilation via arabitol 5-phosphate cannot be excluded since AtlD can also use Ribu5P, albeit with reduced efficiency as shown both by the kinetic parameters (Table 2) and the significant residual Ribu5P detected in the enzyme reactions (Fig. S2). Moreover, *in vitro* enzymatic analyses of purified arabitol phosphate dehydrogenase from *E. avium* (23) and *Bacillus halodurans* (56) showed that both could convert arabitol 1-phosphate and arabitol 5-phosphate into both Xyl5P and Ribu5P. By assessing metabolic operations *in vivo*, ^13^C-MFA experiments should identify which assimilation pathway is actually used *in vivo*.

### 3.2 ^13^C-MFA experimental design

Experimental design is a key step to define the isotopic composition of the label input and the isotopic data to be measured, thereby improving both the number of fluxes that can be estimated from a set of isotopic data and the precision of the flux values. To address this point, we used a dedicated program, IsoDesign (57), using as input metabolic models of each carbon source and several labels as described in the material and methods section. For arabitol and mannitol, good flux precision could be obtained with mass spectrometry labelling data from proteinogenic amino acids. This is advantageous for two reasons. First, the experimental setup is simpler than for intracellular metabolites because no quenching is required and the cellular pellet can just be collected by centrifugation. Second, the labelling can be measured by NMR and MS, providing crucial positional information to distinguish between the two pathways (i.e. via arabitol 5-phosphate or via arabitol 1-phosphate dehydrogenase). Among the different label inputs tested, 100% [5-^13^C] arabitol appeared ideal to identify whether arabitol is converted into arabitol 1-phosphate only or into arabitol 5-phosphate only. In addition, a 9:1 mix of [1-^13^C] and [2-^13^C] arabitol was used to see if both pathways are active. For mannitol, the best label input was 100 % [1-^13^C]. Finally, since methanol is a C1 compound, a stationary ^13^C-MFA approach is not suitable since the amino acids become fully labeled in the isotopic steady-state. Non-stationary ^13^C fluxomics should be used instead to follow the incorporation of the tracer after a pulse of labeled substrate; however significantly more ^13^C data are required for this approach (58, 59) (Supplementary Data 1).

All labeling samples were collected in the exponential growth phase at metabolic steady state and analyzed by MS or 1D ^1^H NMR. The labelling profile of the analyzed metabolites, as measured by their CIDs, were inspected manually and then used with additional NMR and physiological data to fit a model of *B. methanolicus*’s central metabolism (Supplementary Data 1).

### 3.3. Measurement of physiological parameters

Assessment of physiological parameters is a prerequisite for flux calculation. Here, *B. methanolicus* was grown in batch on three different carbon sources. For each cultivation, growth rate and consumption and production rates were determined. Results are given in Table 3. No significant differences between the physiological parameters obtained for the culture replicates were observed. On methanol (batch cultures at 50 °C), around 35 % of the carbon source was directly evaporated, and biomass and carbon dioxide were the only products formed in detectable amounts. The maximal growth rate obtained here with methanol (0.46 h^−1^) is slightly higher than reported in a previous proteomic study of *B. methanolicus* MGA3 (0.40 h^−1^, (14)) or for the growth of the related strain *B. methanolicus* PB1 (0.32 h^−1^, (15)). Biomass yields did not differ significantly from 0.5 g/g indicating that approximately half of the consumed methanol went to biomass and the other half was oxidized to CO_2_. The growth rates with mannitol and arabitol are consistent with published values (22)).

**Table 3:**
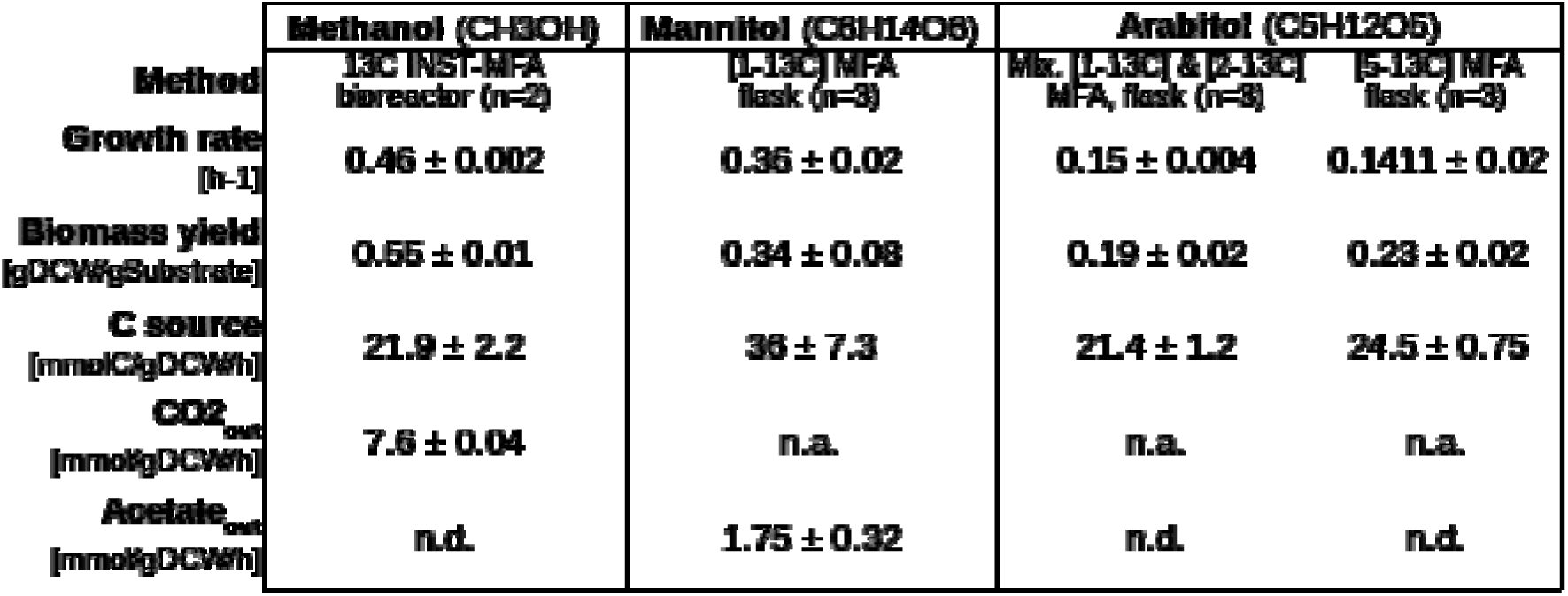
Physiological parameters of *B. methanolicus* cultures used for Metabolic Flux Analysis. Growth rate and biomass quantities were deduced from OD_600_ measurements in the exponential growth phase. Carbon source evolution rates were measured from supernatant samples and analyzed by NMR. Labelled CO_2_ was monitored by MS in the bioreactor’s outgoing gas flow. All cultures were aerobic. n.a.: not measured; n.d.: not detected. The uncertaintie shown are standard errors between biological replicates.

*B. methanolicus* MGA3 is known to overproduce glutamate at up to 50 g/l (21) on methanol under magnesium or methanol limitation (60, 61). When grown on arabitol or methanol, no metabolite accumulated in the supernatant at concentrations above the NMR detection limit (approximately 100 µM). There was therefore little or no metabolite secretion by MGA3 under our chosen methanol and arabitol growth conditions, which is consistent with previous studies and the growth conditions studied here (27, 28). However, acetate was produced at up to 2 mmol/gDCW/h in mannitol cultures (yield, 0.3 mol/mol). To the best of our knowledge, acetate production has never previously been reported for *B. methanolicus*. Based on genomics data (21), we hypothesized that acetate is synthesized from acetyl-CoA in a classical two-step process involving phosphate acetyltransferase (EC:2.3.1.8, BMMGA3_RS15725) and acetate kinase (EC:2.7.2.1, BMMGA3_RS12735). Acetate has moreover been discussed at length in the context of overflow metabolism, a special metabolic state in which fermentation pathways are used even though further oxidation (respiration) would be more ATP-efficient (62–65).

Overall, we show the reproducible growth characteristics of our cultures, and show an unexpected production of acetate that may have an impact for industrial applications, as overflow metabolism leads to carbon and energy waste.

### 3.4 *In vivo* characterization of arabitol assimilation

The *in vitro* enzymatic analyses of the two dehydrogenases, AltD and AltF, suggest that both assimilation pathways (i.e. via arabitol 1-phosphate and via arabitol 5-phosphate) may operate *in vivo*. To confirm whether either or both arabitol assimilation pathways are operative in *B. methanolicus*, as suggested for *E. avium* (23) and *Bacillus halodurans* (56), we carried out a ^13^C-MFA specifically designed to discriminate between the two pathways (see section 3.1, and Fig 2). Interestingly, while the possibility to assimilate arabitol through Xyl5P or Ribu5P or both was a free parameter, the optimal solution found during the fitting process exclusively used the route through Xyl5P (Fig. 1C). This indicates that the (low) activity observed *in vitro* through Ribu5P was not present at a detectable level in our cultures, and that the PTS system imports arabitol as arabitol 1-phosphate. The kinetic parameters obtained for AtlD and AtlF are in line with the flux data, i.e. entry of arabitol into the pentose phosphate pathway (PPP) via PTS-mediated uptake and phosphorylation to arabitol 1-phosphate followed by oxidation to Xyl5P with AtlD as the major dehydrogenase (Fig. 2 and Table 2).

Overall, these data demonstrate for the first time how arabitol is assimilated in *B. methanolicus* and rule out the hypothesis of additional catabolism through arabitol 5-phosphate and Ribu5P derived from our enzymatic analyses (Table 2) and previous reports in *E. avium* (23) and *B. halodurans* (56).

### 3.5. *In vivo* operation of the pentose phosphate pathway with the different carbon sources

*B. methanolicus* assimilates methanol through the ribulose monophosphate (RuMP) cycle that condenses formaldehyde (For) and ribulose 5-P (Ribu5P) into hexulose 6-P (Hex6P) (66). The regenerative part of the RuMP cycle that maintains a pool of Ribu5P overlaps with the non-oxidative pentose phosphate pathway (PPP). Strong flux through the non-oxidative part of the PPP is therefore expected on methanol and indeed, ribulose-phosphate 3-epimerase’s relative flux accounted for 62 % of methanol assimilation (*rpe*, Fig. 1A). Interestingly, mannitol and arabitol are closely connected to the PPP since mannitol is converted to fructose 6-P (F6P) (just like methanol), whereas arabitol is converted to Xyl5P as discussed above. On mannitol and arabitol (Fig. 1B), PPP utilization was lower compare to methanol. However on arabitol, the fluxes associated with the PPP accounted for a larger fraction of the assimilated carbon flow than on mannitol (the estimated *rpe* flux was 15% and 28 % of respectively mannitol and arabitol assimilation; Fig 1B-C). This indicates that the non-oxidative PPP is more important for carbon assimilation on arabitol than on mannitol, in spite of similar expression levels (22). Simulations carried without transaldolase activity indicated that its activity was essential to fit the labeling data given the network topology used (notably *glpx* been irreversible) on arabitol. A possibility for future studies would be to take advantage of adaptive laboratory driven evolution, or overexpression of key enzymes such as transaldolase, to investigate if the PPP could be adjusted to increase growth rates when arabitol is the sole source of carbon and energy.

The RuMP cycle has several variants, which differ in their efficiency (67). Genes for two of these have been identified in MGA3, namely, the fructose bisphosphate aldolase/sedoheptulose bisphosphatase (SBPase) cycle (53, 68) and the fructose bisphosphate aldolase/transaldolase (TA) cycle(69), which as their names suggest favor the regeneration of Ribu5P through sedoheptulose bisphosphatase and transaldolase, respectively. It is generally accepted that MGA3 uses the SBPase variant (66). The main evidence for this is the presence of a copy of a characteristic gene of the SBPase variant (*glpX*^P^) on the pBM19 plasmid, whereas there is only one transaldolase gene (*ta*^C^) in the chromosome. Proteomic (26) and transcriptomic (21) studies have also associated *glpX*^P^ with a significant increase in expression in methanol compared with mannitol, whereas the expression associated with *ta*^C^ remained constant. According to the ^13^C-MFA, both *glpx* (associated to *glpX*^P^) and *ta* (associated to *ta*^C^) carried a comparable flux, thus both variant may be active (Fig. 3). This arrangement may serve as a fail-safe to guarantee the replenishment of Ribu5P. Alternatively, since transaldolase activity is essential to fit the isotopic data for growth on arabitol, an advantage of the TA cycle may be that it increases the flexibility of the PPP and allows the regeneration of important precursors such as Ribu5P from different carbon sources.

**Fig. 3:**
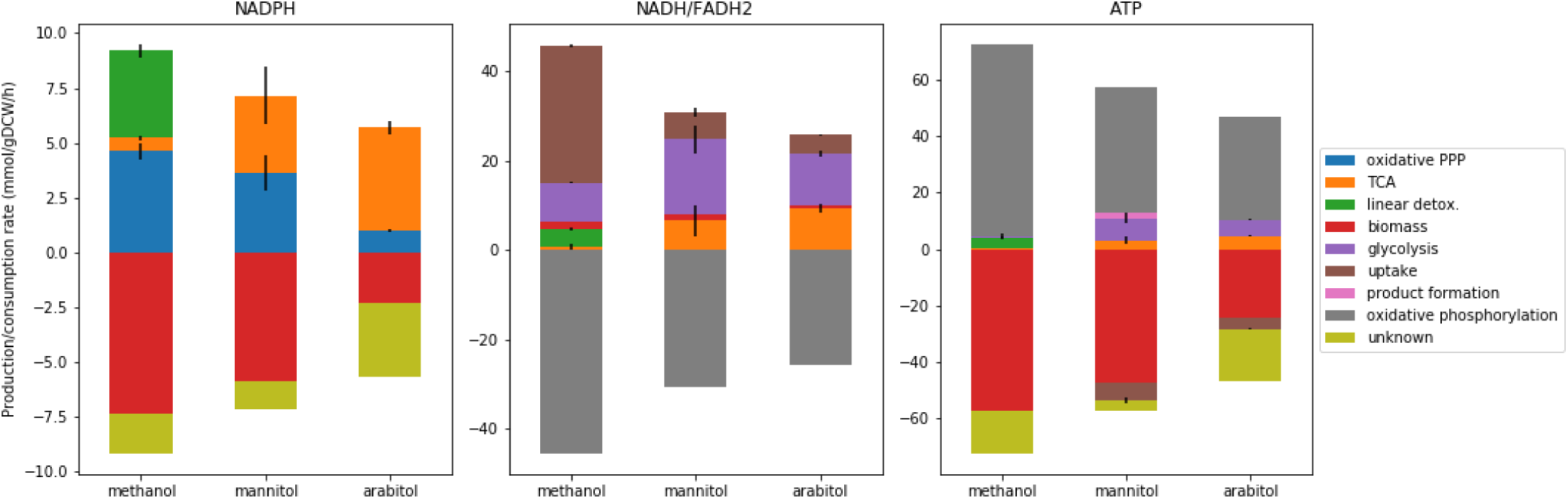
Estimated absolute rates of production and consumption of NADPH, NADH/FADH2, and ATP. Rates (mmol/gDCW/h) are calculated as the sum of the estimated flux of the reactions modeled in the MFA that are expected to produce (positive value) or consume (negative value) those cofactors. Values are averaged over the biological replicates and the error bars are the associated standard deviation (n_methanol_ = 2, n_arabitol_ = n_mannitol_ = 3). The growth requirements (“biomass”, red) are computed from measures on *B. subtilis* (mmol/gDCW) and the growth rates we observed. Production and consumption rates should be balanced in our conditions, so we mark putative production/consumption rates as “unknown” to complete the balance when needed. The full production of NADH and FADH2 is assumed to be consumed in the respiratory chain (“oxidative phosphorylation”) and to produce ATP (with P/O_NADH_ = 1.5, P/O_FADH2_ = 1). We modeled the PTS systems of mannitol and arabitol as consumers of one equivalent ATP (in “uptake”). NADPH production: glucose 6-phosphate dehydrogenase (*zwf*), phosphogluconate dehydrogenase (*gnd*), isocitrate dehydrogenase (*idh*), and methylenetetrahydrofolate dehydrogenase from the linear detoxification pathway (detox). NADH production: glyceraldehyde-3-phosphate dehydrogenase (*pgk*), pyruvate dehydrogenase (*pdh*), 2-oxoglutarate dehydrogenase (*akgdh*), malate dehydrogenase (assimilated to *fum*), methanol, mannitol and arabitol dehydrogenase (*mdh*, manupt, araupt), and formate dehydrogenase from the linear detoxification pathway (detox). FADH2 production: succinate dehydrogenase (*fum*). ATP consumption: 6-phosphofructokinase (*pfk*), PTS systems of mannitol and arabitol (manupt, araupt). ATP production: phosphoglycerate kinase (*pgk*), pyruvate kinase (*pyk*), succinyl-CoA synthetase (assimilated to *akgdh*), acetate kinase (*out_Ac*), and formate-tetrahydrofolate ligase from the linear detoxification pathway (detox).

The labeling data suggest that the oxidative part of the PPP was similar on methanol and on arabitol and almost five times higher on mannitol (the estimated relative glucose 6-phosphate isomerase (pgi) flux was 31, 12 and 9 %, respectively; Fig. 1). This follows the conclusions of previous studies showing that the oxidative part of the PPP is slightly upregulated on mannitol compare to methanol (14) but suggested that this route should be used on methanol to provide NADPH while detoxifying formaldehyde (14, 27) via the so-called cyclic dissimilatory RuMP pathway (21, 70). The utilization of the oxidative part of the PPP on methanol is of particular interest because, intuitively, one imagines that decarboxylation should be avoided when growth occurs on a C1 to avoid wasting carbon. This intuition proved to be correct in Bennett et al.’s (71) study of an *E. coli* synthetic methylotroph in which they knocked-out *pgi* to increase the regeneration of Ribu5P through the non-oxidative part of the PPP. However, we cannot exclude the possibility that the oxidative part of the PPP serves as a backup formaldehyde dissimilation pathway when the linear dissimilation pathways become saturated at high methanol concentrations. *B. methanolicus* is indeed quite sensitive to variations in methanol concentration (72) and we can assume that this critical biological function is tightly controlled.

As expected, the ^13^C-MFA shows that the PPP is critical on methanol as it overlaps with the RuMP responsible for methanol assimilation. The SBPase variant of the RuMP is active, however we could not rule out a parallel operation of the TA variant. We suggest that a parallel operation on mixed carbon sources may benefit *B. methanolicus* to replenish important precursor’s pools.

### 3.6 *In vivo* operation of the TCA cycle with the different carbon sources

*B. methanolicus* has a full gene set for a functional tricarboxylic acid cycle (TCA) and glyoxylate shunt (14, 21). This seems to contrast with some methylotrophs, including some that use the RuMP pathway, that do not need a complete TCA to fulfill their energy requirements (73). Results (Fig. 1A) indicated that the TCA was used much less than the RuMP pathway on methanol, with relative fluxes 3% up to isocitrate (ICit) and almost no flux afterwards (Fig. 1A). This small flux is needed to support the synthesis of biomass precursors. On mannitol and arabitol in contrast, the TCA was used intensively, with relative flux of respectively 57 % and 110% for isocitrate dehydrogenase (*idh*) (Fig.1B-C). These results are in agreement with previous measurements of the actual usage of the TCA during methylotrophic growth. At the transcript and protein levels, it has been suggested that the TCA should be more active on mannitol than on methanol (21, 26). Activity assays in crude cell extracts also showed very low 2-oxoglutarate dehydrogenase (*akgdh*) activity (60). Finally, isotopic labeling experiments have shown slow isotopic enrichment of key TCA metabolites (citrate, 2-oxoglutarate, fumarate) on methanol (27).

Although *B. methanolicus* also has glyoxylate shunt genes (BMMGA3_RS01750, 2.3.3.9; and BMMGA3_RS01755, 4.1.3.1), the reported expression levels suggest that this pathway does not carry a high flux under mannitol or under methanol growth conditions (21, 26). In agreement with these findings, the glyoxylate shunt fluxes estimated in this study were negligible under all the tested conditions (Fig. 1).

Overall, as reported before, the TCA is mainly active in non-methylotrophic conditions and the glyoxylate shunt was inactive in our conditions.

### 3.7 Analysis of cofactor usage with the various carbon sources

To assess how *B. methanolicus* balances its redox and energy needs, we used the estimated fluxes from the ^13^C-MFA to infer production and consumption rates of ATP, NADH and NADPH, as displayed in Fig. 4. ^13^C-MFA is constrained by carbon balance and the distribution of the isotopic tracer, but unlike flux balance analysis it is typically not constrained by cofactor balance. We used the estimated fluxes from the MFA and the expected stoichiometry of the associated reactions to assess the absolute rates of cofactor production and consumption. In the absence of specific measurements, we use the biomass requirements of *B. subtilis* (8, 74). Additionally, since the samples were collected in metabolic pseudo steady-state, the production and consumption rates of each cofactor were assumed to be balanced; this allowed us to identify rates that would have been impossible to estimate otherwise such as the proportion of NAD (and ATP) formed from the respiratory chain (with a P/O=1.5) and other production/consumption rates not accounted for by our other assumptions.

Despite their different growth rates, which influence their cofactor requirements for growth (Table 3), the cells grown on mannitol and arabitol had a similar absolute usage of NADH, NADPH and ATP (Fig. 3). ATP and NADH came mostly from the same sources. However, the proportion of NADPH generated in the oxidative part of the PPP was three times higher in mannitol growth conditions than in arabitol (3.5 against 1.0 mmol/gDCW/h) which is consistent with the higher fluxes observe in the oxidative part of the PPP on mannitol. This difference was compensated by a higher flux in the TCA for arabitol (4.7 against 3.5 mmol/gDCW/h) in excess of the estimated NADPH requirements for growth and even contributing to the production of ∼3.4 mmol/gDCW/h NADPH whose consumption remains unaccounted for. Discrepancies between estimates of the cofactor requirements for biomass formation and those derived from measured isotopic data are typical for this kind of analysis (75, 76), and can be attributed to an underestimation of cofactor requirements, notably for non-essential processes such as cell motility (for ATP).

Comparing methylotrophic and non-methylotrophic growth, it is not surprising that NADH was produced at a higher rate on methanol (45.6 mmol/gDCW/h, versus 30.4 and 25.7 mmol/gDCW/h on mannitol and arabitol respectively), because of the conversion of methanol into formaldehyde by methanol dehydrogenase (77). This suggests that at the same growth rate (i.e. methanol vs mannitol), O_2_ consumption should be higher on methanol to provide the additional NAD^+^ required. The NADPH total production was estimated to be largely higher on methanol (9.2 mmol/gDCW/h) compare to mannitol and arabitol (Fig. 3). It was mainly provided by the oxidative part of the PPP (which would be part of the cyclic dissemination pathway; 4.6 mmol/gDCW/h) as on mannitol but also by the linear formaldehyde detoxification pathways, while usage of the TCA was negligible. These estimates suggest that linear detoxification pathways play an important role in the generation of NADPH on methanol, in addition to their established protective function against formaldehyde.

Overal, non-methylotrophic growth on mannitol and arabitol share the same features despite their associated different growth rates (with the notable exception of the contribution of the oxidative PPP for NADPH production). On methanol, however, we observe more differences in the origin of cofactors production which clearly highlight a different metabolic state. Importantly, the linear detoxification pathways share with the PPP an important role in cofactor regeneration.

### 3.8. Conclusion

In summary, this MFA of *B. methanolicus* MGA3 provides three snapshots of its metabolic states for growth on methanol, mannitol or arabitol. Isotopic data are consistent with prior knowledge of MGA3 methylotrophy, showing greater flux in the RuMP cycle than in the TCA. The ^13^C-MFA provided new insights related to the utilization of cyclic RuMP versus linear dissimilation pathways, and between the RuMP cycle variants; and finally, the characterization of the arabitol assimilation pathway was completed using enzymatic data. In future studies, these validated flux maps will be used as references for constraint based modelling to validate genome-scale model predictions. Overall, the information provided in this work and previous omics studies on *B. methanolicus* metabolism can be used to improve design strategies for new strains (i.e. by multiaomics analysis). For instance, the comparison between methanol and arabitol might help identifying key control steps in the metabolic network that allows shifting from a cyclic (i.e. methanol with the Rump cycle) to a linear (i.e. arabitol with the PPP) mode of operation. The experimental path outlined here likely leads to *B. methanolicus* becoming a viable alternative to existing cell factories.

## ACKNOWLEDGEMENT

MetaToul (www.metatoul.fr), which is part of MetaboHub (ANR-11-INBS-0010, www.metabohub.fr) are gratefully acknowledged for their help in collecting, processing and interpreting 13-C NMR and MS data. We would like to thank Marcus Persicke, Maud Heuillet, Edern Cahoreau, Lindsay Periga, Pierre Millard, Gilles Vieira and Jean-Charles Portais for the insights they provided.

## ABBREVIATIONS

13C-MFA: 13C metabolic flux analysis
CID: carbon isotopologue distribution
FBA: Flux Balance Analysis
MS: mass spectrometry
NMR: nuclear magnetic resonance
PTS: phosphotransferase system

**Pathways**
SBPase: fructose bisphosphate aldolase/sedoheptulose bisphosphatase
TA: fructose bisphosphate aldolase/transaldolase
PPP: pentose phosphate pathway
RuMP: Ribulose Monophosphate Cycle
TCA: Krebs cycle

**Reactions**
akgdh: 2-oxoglutarate dehydrogenase and 2-oxoglutarate synthase
Araupt: arabitol uptake
detox: linear detoxification pathways
fum: fumarate reductase
glpx: sedoheptulose-bisphosphatase
gnd: phosphogluconate dehydrogenase
hps: 3-hexulose-6-phosphate synthase
idh: isocitrate dehydrogenase
Manupt: mannitol uptake
mdh: methanol dehydrogenase
pdh: pyruvate dehydrogenase
pfk: 6-phosphofructokinase and fructose-bisphosphatase
pgi: glucose 6-phosphate isomerase
pgk: glyceraldehyde 3-phosphate dehydrogenase, phosphoglycerate kinase, and phosphoglycerate mutase
phi: 6-phospho-3-hexuloisomerase
pyk: pyruvate kinase
rpe: ribulose-phosphate 3-epimerase
rpi: ribose 5-phosphate isomerase
rpi: ribulose phosphate isomerase
ta: transaldolase
tkt1, tkt2: transketolases
zwf: glucose-6-phosphate dehydrogenase and 6-phosphogluconolactonase.

**Metabolites**
Ace: acetate
Aco: aconitate
AKG: 2-oxoglutarate
ala: alanine
Ara: arabitol
arg: arginine
asp: aspartate
Cit: citrate
DHAP: dihydroxyacetone phosphate
Ery4P: erythrose 4-phosphate
For: formaldehyde
Fru6P: fructose 6-phosphate
FruBP: fructose 1,6-bisphosphate
Fum: fumarate
G3P: 3-phospho-D-glycerate
GAP: glyceraldehyde 3-phosphate
Glc6P: glucose 6-phosphate
glu: glutamate
gly: glycine
GlyOx: glyoxylate
Gnt6P: 6-phosphogluconate
Hex6P: hexulose 6-phosphate
his: histidine
ile: isoleucine
leu: leucine
lys: lysine
Mal: malate
Man: mannitol
MeOH: methanol
met: methionine
OAA: oxaloacetate
PEP: phosphoenolpyruvate
PGA: 2-phospho-D-glycerate
phe: phenylalanine
pro: proline
Pyr: pyruvate
Rib5P: ribose 5-phosphate
Ribu5P: ribulose 5-phosphate
Sed7P: sedoheptulose 7-phosphate
ser: serine
Suc: succinate
thr: threonine
tyr: tyrosine
val: valine
Xyl5P: xylulose 5-phosphate.

