## Supplementary Text and Figures for "Charting the metabolic landscape of the facultative methylotroph *Bacillus methanolicus*"

### Text S1: Phylogeny of AltD and AltF

AtlD was found to be the major dehydrogenase preferring Xyl5P over Ribu5P and NADH over NADPH as cofactor (Table 2). The *V*_max_ of AtlD with Xyl5P and NADH of 1.33 ± 0.23 U mg^-1^ was around 10-fold lower than the values reported for the *E. avium* arabitol phosphate dehydrogenase (APDH) (1) and around 20-fold lower than those reported for the *B. halodurans* APDH (2), however, affinity for Xyl5P as substrate is 3-fold higher with a *K*_M_ value of 0.07 ± 0.03 mM in both cases. With Ribu5P as substrate, AtlD showed a *V*_max_ of around 37 % of that observed with Xyl5P, which is higher as compared to the 2 to 3 % reported in *E. avium* (1). With a value of 0.49 ± 0.06 U mg^-1^ the *V*_max_ of AtlD with Ribu5P as substrate is around 7-fold lower than what was observed in *B. halodurans* (2), however, affinity for Ribu5P is around 3-fold higher in the case of AtlD. Comparison of the *E. avium* and *B. halodurans* APDHs to AtlD reveals that the three enzymes have similar kinetic characteristics: they all prefer Xyl5P over Ribu5P and subsequently show increased *V*_max_ values with the former; however, the *B. halodurans* APDH was reported to have a wider substrate specificity since it can also reduce fructose 5-phosphate and ribose 5-phosphate (2).

It was recently found that *Listeria monocytogenes* possesses three pentitol phosphate dehydrogenases showing arabitol phosphate dehydrogenase activity: Lmo2663, Lmo2664 and Lmo0506 (3). Lmo2663 and Lmo2664 are encoded in the same operon and were shown to reduce Xyl5P, while only Lmo2664 also reduced Ribu5P, but at a much lower rate. The reduction rate of Xyl5P is 3- and 5-fold higher for Lmo2663 and Lmo2664, respectively, than for AtlD, while reduction of Ribu5P is around 8-fold higher for AtlD than Lmo2664. In regard to Lmo0506, AtlD showed 7-fold higher Xyl5P and Ribu5P reduction rates, however, the product of the reduction of Ribu5P by Lmo0506 is expected to yield ribitol 5-phosphate and not arabitol 5-phosphate, even when ribitol could not be utilized by *L. monocytogenes* (3). Lmo2663 and Lmo2664 seem to be the major arabitol phosphate dehydrogenases in *L. monocytogenes*, with Lmo2664 showing a higher efficiency and resemblance to kinetic characteristics shown for the *E. avium* (1) and the *B. halodurans* (2) APDHs. Even though Lmo0506 exhibits strong sequence similarity to Lmo2663 and Lmo2664, activities were significantly lower and more in the range of those measured for AtlF: 0.19 U mg^-1^ (3) vs. 0.11 ± 0.01 U mg^-1^, respectively.

In all tested reactions NADH was the preferred cofactor for AtlD with a *K*_M_ value of 0.01 ± 0.01 mM, which was 10-fold lower than the determined *K*_M_ for NADPH and equivalent to data previously reported for *E. avium* (1) and *B. halodurans* (2). Interestingly, neither Lmo2663 nor Lmo2664 from *L. monocytogenes* accepted NADPH as a reducing substrate (3).

The closest matches shown for AtlD and AtlF when subjected to BLASTp analysis are proteins annotated as sorbitol dehydrogenases and galactitol phosphate dehydrogenases, respectively. To our knowledge and except for the here presented AtlD and AtlF, the *E. avium* (1), *B. halodurans* (2) and *L. monocytogenes* (3) proteins, none of those enzymes have been characterized. More precisely, characterization of the *B. halodurans* APDH derived from an effort to verify that high-scoring homologues shared substrate specificity with the APDH of *E. avium* (1), suggesting that other uncharacterized proteins with high homology might encode arabitol phosphate dehydrogenases. Additionally, several polyol dehydrogenases have been characterized for *Rhodobacter sphaeroides* which show substrate specificity to more than one substrate (4, 5). More precisely, it was shown that both sorbitol and mannitol dehydrogenases could accept galactitol as non-phosphorylated substrate, even when *R. sphaeroides* was unable to grow with galactitol as sole source of carbon (6) and characterization of the galactitol dehydrogenase showed that it accepted several non-phosphorylated alcohols that presented different binding modes within the substrate binding pocket (5).

Homology comparisons of the AtlD and AtlF sequences place them inside the superfamily of medium-chain dehydrogenases/reductases (MDRs) (accession: cl16912), although each belong to a different sub-family based upon sequence similarities (Fig. S4). MDRs display a broad range of activities with several members with unknown functions to date. The total number of families is estimated to be around 500, with less than 30 % sequence identity between them (7). Indeed, AtlD and AtlF share an amino acid identity at the level of 32 %, much lower than that shared with their respective high-scoring homologues. The complexity and spread of the MDR superfamily defy traditional means of classification, although conservation within the families can be analysed based on mappings to their closest available structures (8). MDRs have a length of around 350 amino acids and generally consist of two domains: a C-terminal coenzyme-binding Rossmann fold domain (PfamA:PF00107) and an N-terminal substrate-binding catalytic domain with distant homology to the GroES structure (PfamA:PF08240) (8, 9). Sequence similarity places AtlD inside the alcohol dehydrogenase family of the MDR superfamily that contains a putative NAD(P) binding site and both a catalytic and structural Zn^2+^ binding sites (accession: cd08258), while AtlF falls into the sugar dehydrogenases family with a putative NAD(P) binding site and a catalytic Zn^2+^ binding site alone (accession: cd08236). The active site zinc has a catalytic role, while the structural zinc aids in stability. Of the current 86 MDR families characterised, 35 can bind two Zn^2+^ ions per subunit, one catalytic and one structural, 7 can bind one and 38 bind no Zn^2+^ atom, with 54 % of the MDR forms among bacteria belonging to the latter group (8). Moreover, some MDR families consist of members that can either bind two or one zinc atoms, where the latter members have commonly lost the ligands for the structural zinc (8).

According to the aforementioned BLASTp results and the most recent classifications of the MDR superfamily members presented in (7) and (8), we suggest that AtlD belongs to the MDR001 - ADH (alcohol dehydrogenase) family, being the largest MDR family with over 2000 members as of 2010 and containing the classical alcohol dehydrogenases. Comparatively, AtlF would belong to the MDR005 - PDH (polyol dehydrogenase) family (or more precisely to the MDR080 - bPDH (bacterial polyol dehydrogenases) family), containing sorbitol/xylitol dehydrogenases and ᴅ-xylulose reductases but being also able to act on related sugar alcohol substrates. In accordance to what has been mentioned before, ADHs generally have both the structural and catalytic zinc with a catalytic domain conservation of 89 %, being twice of the cofactor binding domain conservation and showing that the catalytic machinery is highly conserved throughout this enzyme family. On the other hand, many sorbitol dehydrogenases of the PDH family bind only the catalytic zinc, especially the ones of bacterial origin. MDR members generally form homodimers, but both bacterial ADHs and PDHs are typically active as tetramers, trait that was experimentally shown for the *E. avium* (1) and the *B. halodurans* (2) APDHs. In addition, there is a recurrence between the number of zinc binding sites and the cofactor preference, with most of the MDR families that bind 2 zinc atoms preferring NAD as cofactor and thus generally acting as dehydrogenases (8).

In order to investigate the interrelations between different MDR members, a phylogenetic tree was built (Fig. S4). While our primary focus lays on the aforementioned characterized arabitol phosphate dehydrogenases, strong matches found by sequence homology searches are also displayed, where bacterial sequences are expectedly predominant. Representative sequences of all three domains of life with a focus on either characterized proteins or model organisms are presented. The phylogenetic tree accuracy is supported by its consistently high bootstrap support values.

We have grouped the major clades and labeled them according to their characterized functions and/or the MDR classification in (7) and (8). Bacterial PDHs and ADHs are shown to be distantly related and each form a common evolutionary branch. Similarly, eukaryotic PDHs and archaeal and Gram-negative TDHs (threonine dehydrogenases) are clustered separately. It is interesting to see that both in the bacterial ADH and PDH groups the members separate in two main evolutionary branches (Fig. S4). If we first focus on the characterized arabitol phosphate dehydrogenases inside the ADH group, we can see that AtlD and *B. halodurans* APDH belong to a different clade than *E. avium* APDH and Lmo2663. Similarly, AtlF falls into a clade with the closest characterized arabitol phosphate dehydrogenase being Lmo0506 and the more distantly related Lmo2664 branches separately. This divergence in both groups could be due the fact that *E. avium* APDH, Lmo2663 and Lmo2664 were shown to prefer Mn^2+^ instead of Zn^2+^ for enzymatic activity or even being inactivated by Zn^2+^ as in the case of *E. avium* APDH (1, 3). Since the effect of metal ions was not tested in AtlD, AtlF or *B. halodurans* APDH, it is only speculative to assume that their respective evolutionary distance to the *E. avium* APDH, Lmo2663 and Lmo2664 indicates that they may belong to the widespread group of MDRs that would require Zn^2+^. Furthermore, Lmo0506 was shown to prefer Zn^2+^ over Mn^2+^, in contrast to the other characterized arabitol phosphate dehydrogenases, and shares the highest sequence homology and is the most closely related to AtlF (Fig. S4). It is somehow unexpected to see the distant relation between the *E. avium* APDH and Lmo2664, and the AtlD and *B. halodurans* APDH, since their mechanisms were shown to be very similar, being all able to reduce both Xyl5P and Ribu5P as substrates. On the other hand, Lmo2663 was shown not to be able to reduce Ribu5P but still shares high sequence homology and evolutionary proximity to APDH (Fig. S4).

The three-dimensional conservation of characterized members within the MDR superfamily points to the existence of ancestral dehydrogenases, which after numerous gene duplicatory events gave rise to the present system of subfamilies (7). Even though classification of MDRs continues to be challenging due to their versatility, their presence in a wide range of life forms strongly indicates the importance of their function.

### Supplementary figures

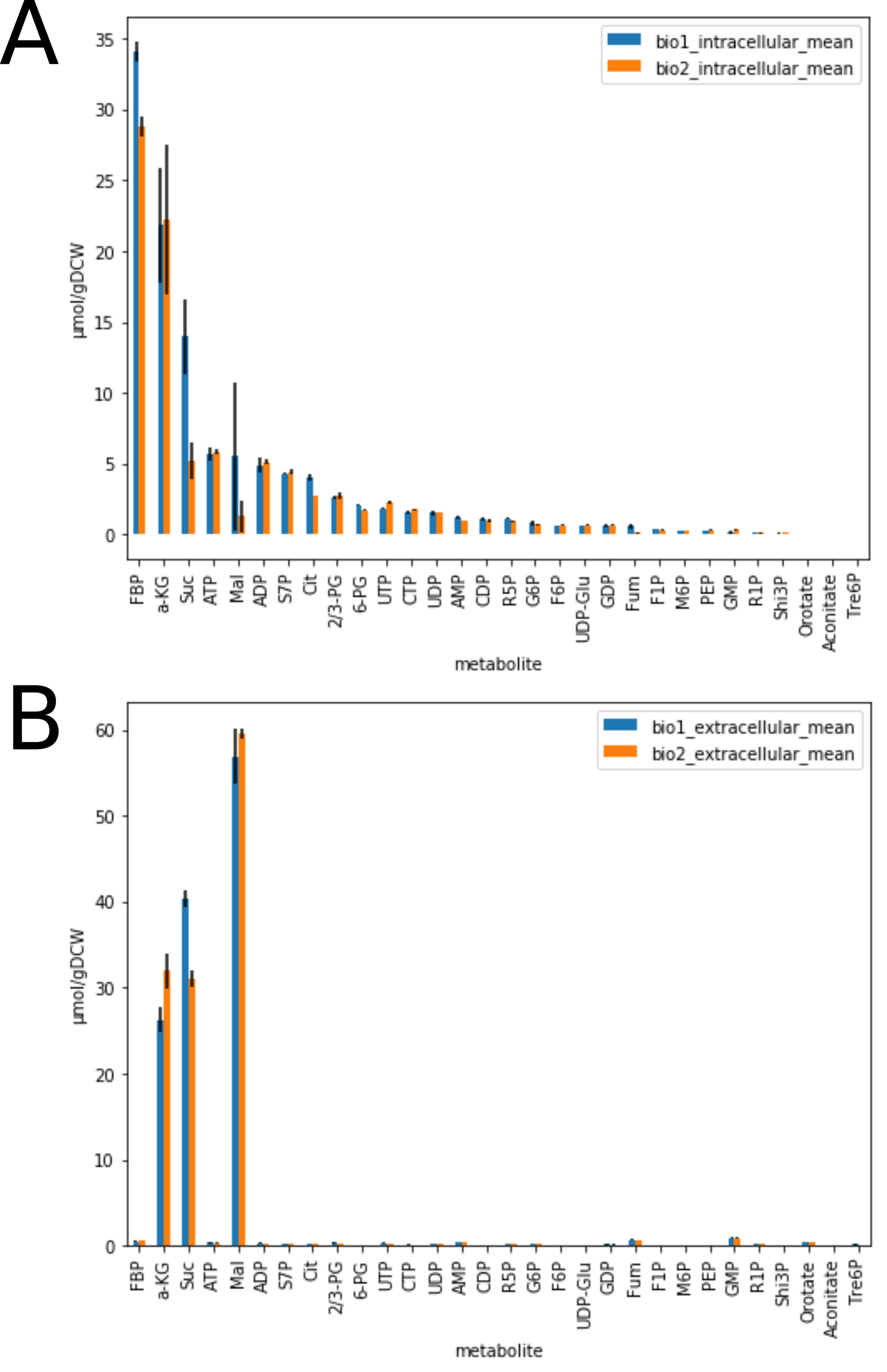

**Fig. S1: Measured intra (A) and extracellular (B) pools of key metabolites of the central metabolism of *B. methanolicus* grown on methanol.** Pools were measured by the “differential method” described in (10). Quickly, this method estimates the intracellular and extracellular metabolites pools from the quantification by ion chromatography mass spectrometry of the metabolites in the whole broth (intra + extracellular) and culture filtrates (extracellular). Samples were collected before the pulse of labeled methanol. The error bars represent the standard deviation of four technical replicates; the two biological replicates are represented by different colors. Metabolites measured were sedoheptulose 7-phosphate (S7P), phosphoenolpyruvate (PEP), 2 + 3 phosphoglycerate (2/3 PG), ribose 5-phosphate + ribulose 5-phosphate + xylulose 5-phosphate (R5P), ribose 1-phosphate (R1P), glucose 6-phosphate (G6P), fructose 1-phosphate (F1P), fructose 6-phosphate (F6P), mannose 6-phosphate (M6P), 6-phosphogluconate (6PG), fructose 1,6-bisphosphate (FBP), adenosine monophosphate (AMP), adenosine diphosphate (ADP), adenosine triphosphate (ATP), UDP-Glucose (UDP-Glc), shikimate 3P (Shi3P), fumarate (Fum), malate (Mal), orotate, citric acid (Cit), oxoglutarate (AKG), succinate (Suc), Trehalose 6-phosphate (Tre6P), Aconitate (Aco), guanosine monophosphate (GMP), guanosine diphosphate (GDP), uridine diphosphate (UDP), cytidine diphosphate (CDP), uridine triphosphate (UTP) and cytidine triphosphate (CTP).

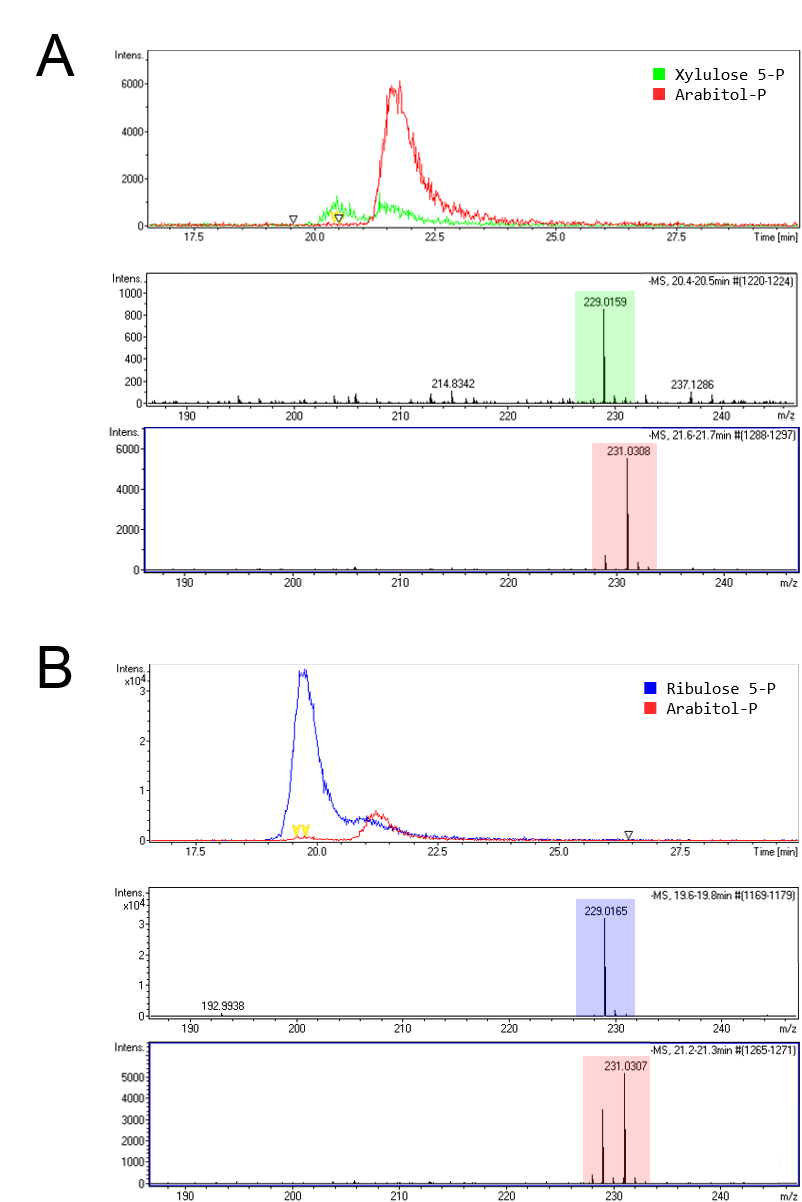

**Fig. S2. Detection of formed arabitol phosphate in AtlD-catalyzed enzyme reactions using LC-MS.** Chromatograms of detected pentitol phosphates (top) and their respective LC-MS spectra (bottom) of enzyme reactions using Xyl5P (A) and Ribu5P (B) as substrates. The peaks for the given compounds were identified by characteristic mass spectra and retention time using standards for Xyl5P and Ribu5P or by comparison to previously reported m/z values for arabitol 5-phosphate (11).

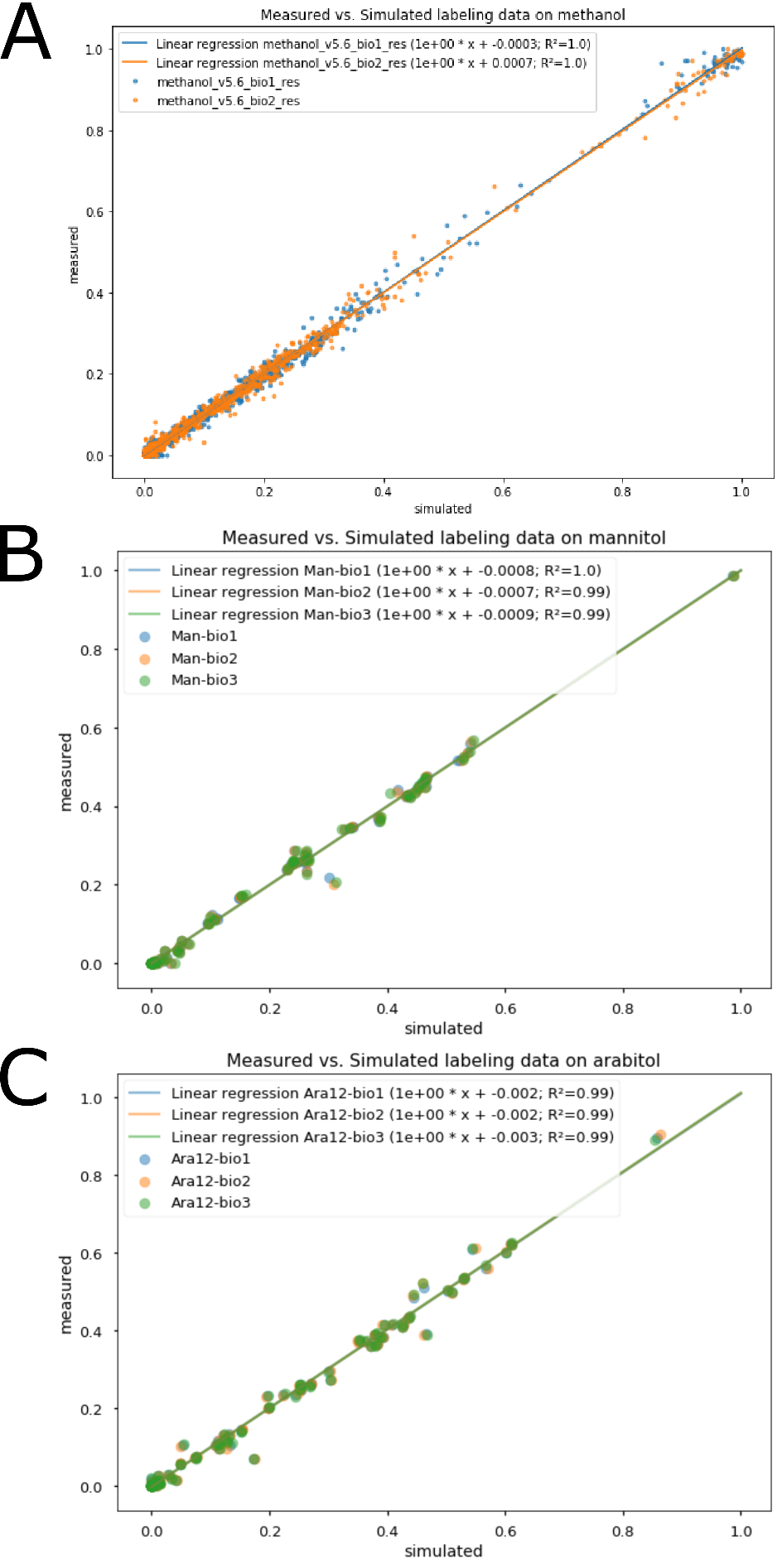

**Fig. S3: Measured vs. estimated labeling proportion in methanol (A), mannitol (B), arabitol (C) conditions.** Each data point refers to the proportion of one of the isotopologues from the Carbon Isotopologue Distribution (CID) of a metabolite used to constrain the models, i.e. either metabolites from central metabolism (A) or proteinogenic amino-acids (B, C) as per described in Methods. On the y-axis, the measured value by mass spectrometry; on the x-axis, the value simulated and used by influx after the optimization process to estimate the fluxes. There is a strong linear correlation between simulated and measured values, which is a good visual indicator of the coherence of the model with experimental data.

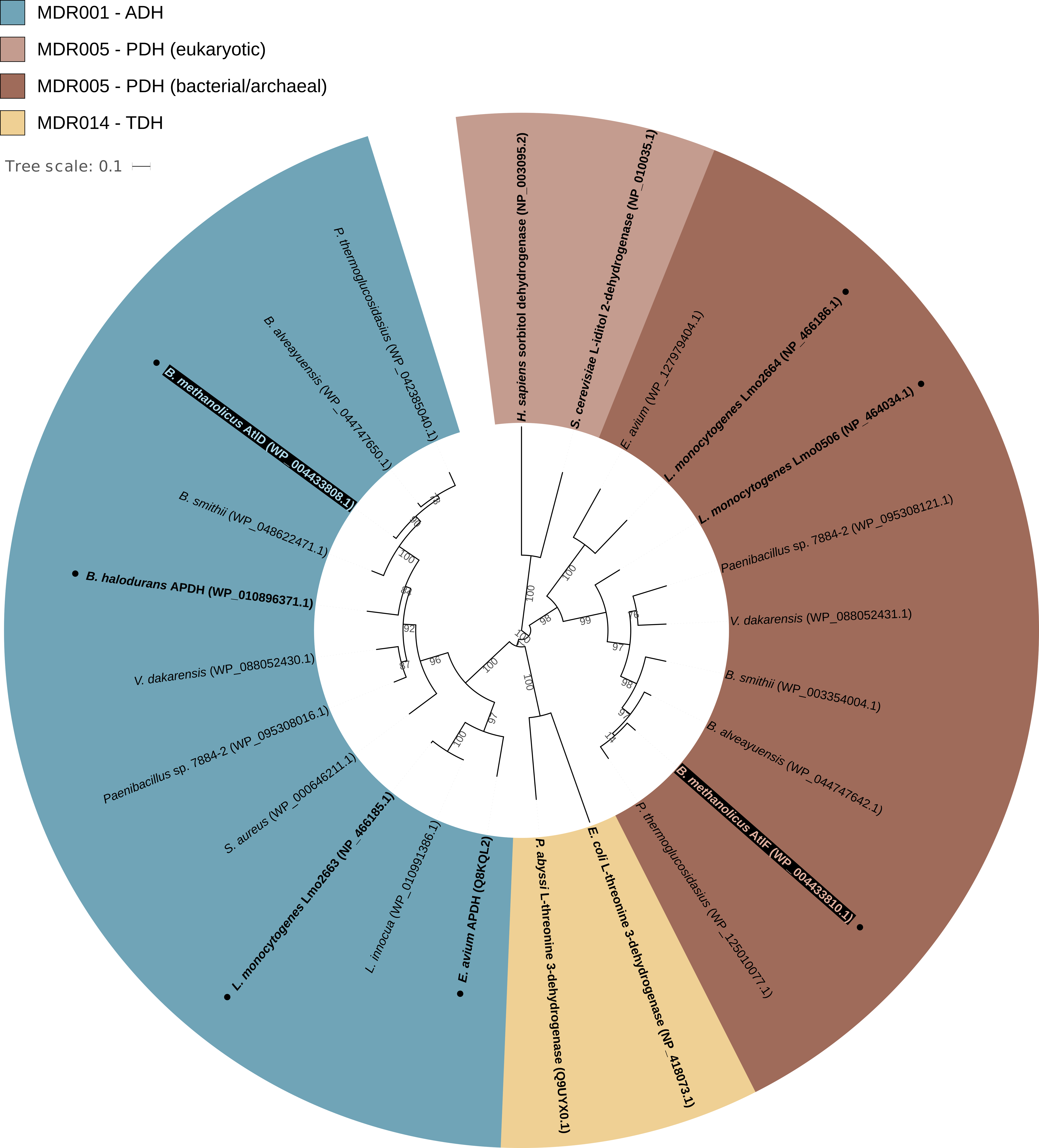
**Fig. S4: Phylogenetic tree of selected medium-chain dehydrogenases/reductase (MDR) homologues.** The tree includes 10 Gram-positive bacterial species, Gram-negative (*Escherichia coli*), archaeal (*Pyrococcus abyssi*), yeast (*Saccharomyces cerevisiae)* and human (*Homo sapiens*) representatives. The suggested member association within each MDR family is highlighted in different colors as indicated in the figure legend. Characterized proteins are shown in bold. Proteins with described arabitol phosphate dehydrogenase activity are highlighted with black dots. Bootstrap support values are indicated as percentage next to the branches. This tree was generated using the Phylogeny Analysis tool (12) and edited using the Interactive Tree of Life online interface (13).
